# A surface pocket in the cytoplasmic domain of the herpes simplex virus fusogen gB controls membrane fusion

**DOI:** 10.1101/2022.03.14.484201

**Authors:** Zemplen Pataki, Erin K. Sanders, Ekaterina E. Heldwein

## Abstract

Membrane fusion during the entry of herpesviruses is carried out by the viral fusogen gB that is activated by its partner protein gH. Unusually, the fusogenic activity of gB is controlled by its cytoplasmic (or intraviral) domain (gB_CTD_) and, according to the current model, the gB_CTD_ is a trimeric, inhibitory clamp that restrains gB in the prefusion conformation. But how the gB_CTD_ clamp is released by gH is unclear. Here, we identified two new regulatory elements within gB and gH from the prototypical herpes simplex virus 1: a surface pocket within the gB_CTD_ and residue V831 within the gH cytoplasmic tail. Mutagenesis and structural modeling suggest that gH V831 interacts with the gB pocket. The gB pocket is located above the “fault line” between adjacent protomers, and we hypothesize that insertion of the gH V831 wedge into the pocket serves to push the protomers apart. This releases the inhibitory clamp on the gB prefusion conformation and activates gB fusogenic activity. Both gB and gH are conserved across all herpesviruses, and this activation mechanism could be used by other gB homologs. Our proposed mechanism emphasizes a central role for the cytoplasmic regions in regulating the activity of a viral fusogen.

**AUTHOR SUMMARY:** Herpes simplex virus 1 (HSV-1) establishes lifelong infections in over a half of people and causes diseases ranging from oral or genital sores to blindness and brain inflammation. No vaccines or curative treatments are currently available. To infect cells, HSV-1 must first penetrate them by merging its lipid envelope with the membrane of the target cell. This process requires the collective actions of several viral and cellular proteins, notably, viral glycoproteins B and H (gB and gH). gH is thought to activate the fusogenic function of gB, but how the two proteins interact is unclear. Here, using mutational analysis, we have identified two new functional elements within the cytoplasmic regions of gB and gH: a surface pocket in gB and a single residue in gH, both of which are important for membrane fusion. Based on structural modeling, we propose that the gB pocket is the binding site for the gH residue, and that their interaction activates gB to cause membrane fusion. These findings extend our knowledge of the HSV-1 membrane fusion mechanism. Mechanistic understanding of HSV-1 entry is essential for understanding its pathogenesis and developing new strategies to prevent infections.

## INTRODUCTION

Membrane fusion during the entry of enveloped viruses is carried out by viral fusogens, which are proteins displayed on the viral surface that bring the opposing viral and host membranes so close that they merge. To do so, these proteins must refold from the high-energy prefusion conformation into the low-energy postfusion conformation. The energy released upon refolding is thought to overcome the large kinetic barrier associated with membrane fusion (reviewed in [1]). To ensure proper spatial and temporal deployment of viral fusogens, their activity is regulated by environmental signals, such as proton concentration, or interactions with other viral and cellular proteins.

Some of the most complex membrane fusion mechanisms are found in herpesviruses – a family of double-stranded-DNA, enveloped viruses that infect most animal species for life. Their entry requires, at a minimum, three conserved glycoproteins, gB, gH, and gL. gB is a transmembrane glycoprotein composed of an ectodomain, a transmembrane helix, and a cytoplasmic domain that functions as a membrane fusogen. By analogy with other viral fusogens, the refolding of gB from the prefusion to the postfusion conformation is thought to provide the energy for membrane fusion. Indeed, the structures of the prefusion [2, 3] and the postfusion forms [4–8] of gB from several herpesviruses suggest large conformational changes that accompany refolding.

gB is a class III fusogen, along with the VSV G, baculovirus gp64, and thogotovirus Gp (reviewed in [9]). Like other class III fusogens, gB exists as a trimer. Yet, uniquely, gB is not a stand-alone fusogen activated by exposure to low pH. Instead, gB must be activated by the conserved heterodimeric complex composed of two viral glycoproteins, gH and gL. gH is a transmembrane glycoprotein composed of an ectodomain, a transmembrane helix, and a short cytoplasmic tail. gL is a soluble glycoprotein that binds gH and is required for its proper folding, trafficking to the cell surface, and function [10, 11].

The gH/gL heterodimer occupies a central place in the herpesvirus entry and membrane fusion processes because it interacts with several key participants. On the one hand, gH/gL interacts with the host cell receptors, either directly or by engaging the viral receptor-binding accessory proteins (reviewed in [12, 13]). On the other hand, it binds and activates gB, the fusogen (reviewed in [12, 13]). According to the prevalent model [14], interaction with the host cell receptor triggers a cascade of events in which gH/gL transmits the activating signal from the host cell receptor to gB. For example, in the prototypical herpesvirus herpes simplex virus 1 (HSV-1) (reviewed in [15]), which establishes lifelong infections in over a half of people ([16] and reviewed in [17]) and causes oral or genital sores (reviewed in [18]) as well as encephalitis (reviewed in [19–21]), binding of the viral receptor-binding protein, gD, to one of its cognate cellular receptors – nectin-1, herpesvirus entry mediator (HVEM), or 3-OS-modified heparan sulfate ([22, 23] and reviewed in [24]) – causes conformational changes in gD that enable it to activate the gH/gL complex [25, 26] that, in turn, activates the fusogenic activity of gB [2, 5, 14, 27] **(Fig. 1)**. But how gB is activated by gH/gL is unknown.

**Figure 1.**
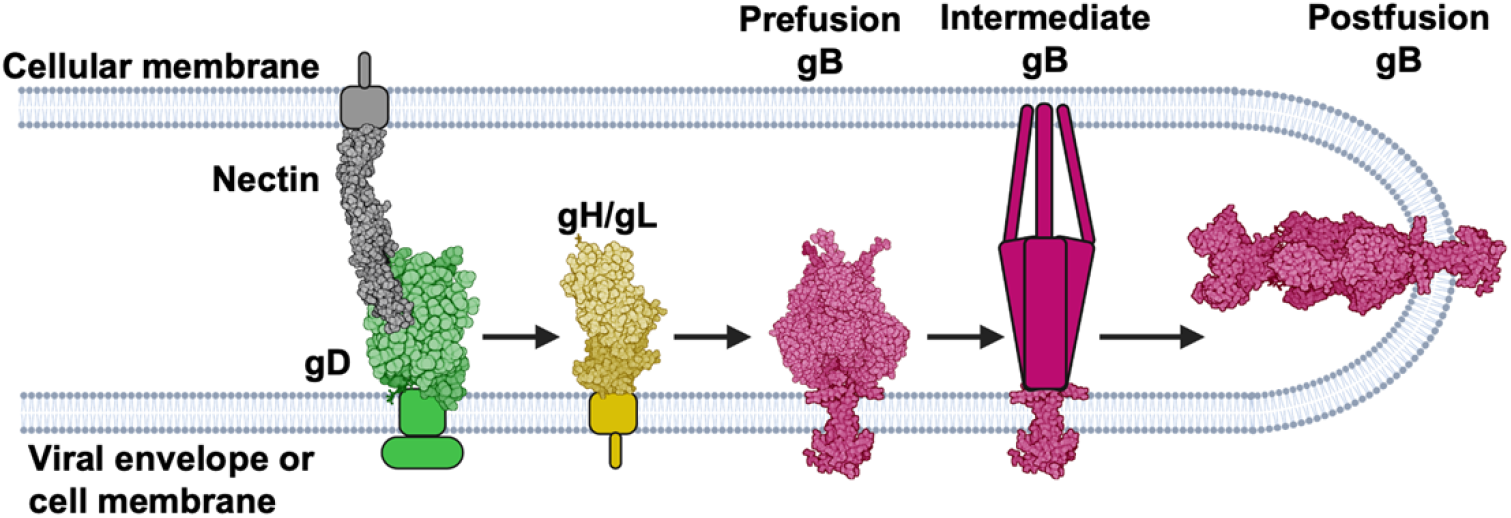
HSV-1 fusion pathway model. gD (2C36 [28]) binds a receptor (3U83 [29]) on the target cell and activates gH/gL; gH/gL (3M1C [27]) triggers gB (6Z9M [2] and 5V2S [5]) to refold and cause fusion. gD has been suggested to be a dimer [28] but is shown here as a monomer for clarity. Figure created with BioRender.com.

While the gB ectodomain undergoes refolding and interacts with the target cell membrane [30], the cytoplasmic domain of gB (gB_CTD_) is also important for fusion. The gB_CTD_ is thought to inhibit the fusogenic activity of gB because most known gB_CTD_ mutations – C-terminal truncations, point mutations, and insertions – are hyperfusogenic, i.e., they increase fusion [31–45]. The crystal structure of the full-length HSV-1 gB [5] revealed that the gB_CTD_ is a trimer stabilized by multiple protein/protein and protein/membrane interactions, and that the majority of the hyperfusogenic gB_CTD_ mutations would be predicted to disrupt these stabilizing interactions [5]. Therefore, we have previously proposed that the gB_CTD_ acts as an inhibitory clamp that stabilizes gB in its prefusion form by restricting conformational rearrangements of the gB ectodomain.

In addition to gB_CTD_, the cytoplasmic tail of gH (gH_CT_) is also important for fusion, but instead of inhibiting fusion, it activates it. gH_CT_ truncations decrease fusion in a manner proportional to the length of the truncated sequence [42] such that the shorter the remaining gH_CT_ length, the lower the fusion. Given the apparent inhibitory function of the gB_CTD_ and the activating function of the gH_CT_, previously, we hypothesized that gH may activate gB by using gH_CT_ to bind the inhibitory gB_CTD_ clamp and disrupt it in a wedge-like manner [5]. However, the respective binding sites on gB_CTD_ and gH_CT_ are unknown.

Here, we identified a previously uncharacterized functional site important for fusion within the HSV-1 gB_CTD_, composed of a surface pocket between adjacent protomers. Mutations of residues A851 and T814 located at the bottom of a surface pocket reduce fusion whereas mutations of residues lining the pocket rim are hyperfusogenic, which suggests that the pocket and the rim have important yet opposite roles in fusion. Moreover, we identified gH_CT_ residue V831 as the most functionally important residue within the gH_CT_. When the gH_CT_ is modelled as an extended polypeptide, V831 ends up approximately the same distance from the membrane as the gB_CTD_ pocket, making interactions at these respective sites plausible if gH and gB were to come into proximity. We hypothesize that gH_CT_ residue V831 serves as the wedge that inserts into the newly identified gB_CTD_ pocket. The gB pocket is located above the “fault line” between adjacent protomers, and we hypothesize that insertion of the gH V831 wedge into the pocket serves to push the protomers apart, which releases the inhibitory clamp. This action would destabilize the inhibitory gB_CTD_ clamp, causing it to release its hold on the gB ectodomain. We hypothesize that in this manner, gH activates the fusogenic activity of gB. The proposed gH-gB triggering mechanism extends our understanding of the regulatory cascade that coordinates HSV-1 entry and may inform new therapeutic approaches aiming at blocking HSV-1 glycoprotein interactions. Both gB and gH are conserved across all herpesviruses, and this activation mechanism could be used by other gB homologs. Our proposed mechanism emphasizes a central role for the cytoplasmic regions in regulating the activity of a viral fusogen.

## RESULTS

### The A851V gB_CTD_ mutant is hypofusogenic

Unlike the very common hyperfusogenic mutations, hypofusogenic gB_CTD_ mutations, i.e., those that decrease fusion, are rare in HSV-1 and HSV-2 and all result in very low surface expression levels, implying a defect in protein folding [32]. Indeed, within the structure of the trimeric HSV-1 gB_CTD_, these mutations map to the hydrophobic core and have been proposed to cause misfolding by eliminating interactions critical for basal trimer stability [5]. No “true” hypofusogenic mutations in the HSV-1 gB_CTD_ that decrease fusion without affecting protein expression have yet been reported. Interestingly, the HSV-1 gB_CTD_ mutant A851V decreased viral entry [31]. Importantly, we found that the A851V mutation had no effect on cell surface expression, as measured by flow cytometry **(Fig. 2a)**. Therefore, we hypothesized that A851V reduces viral entry by reducing gB fusogenicity.

**Figure 2.**
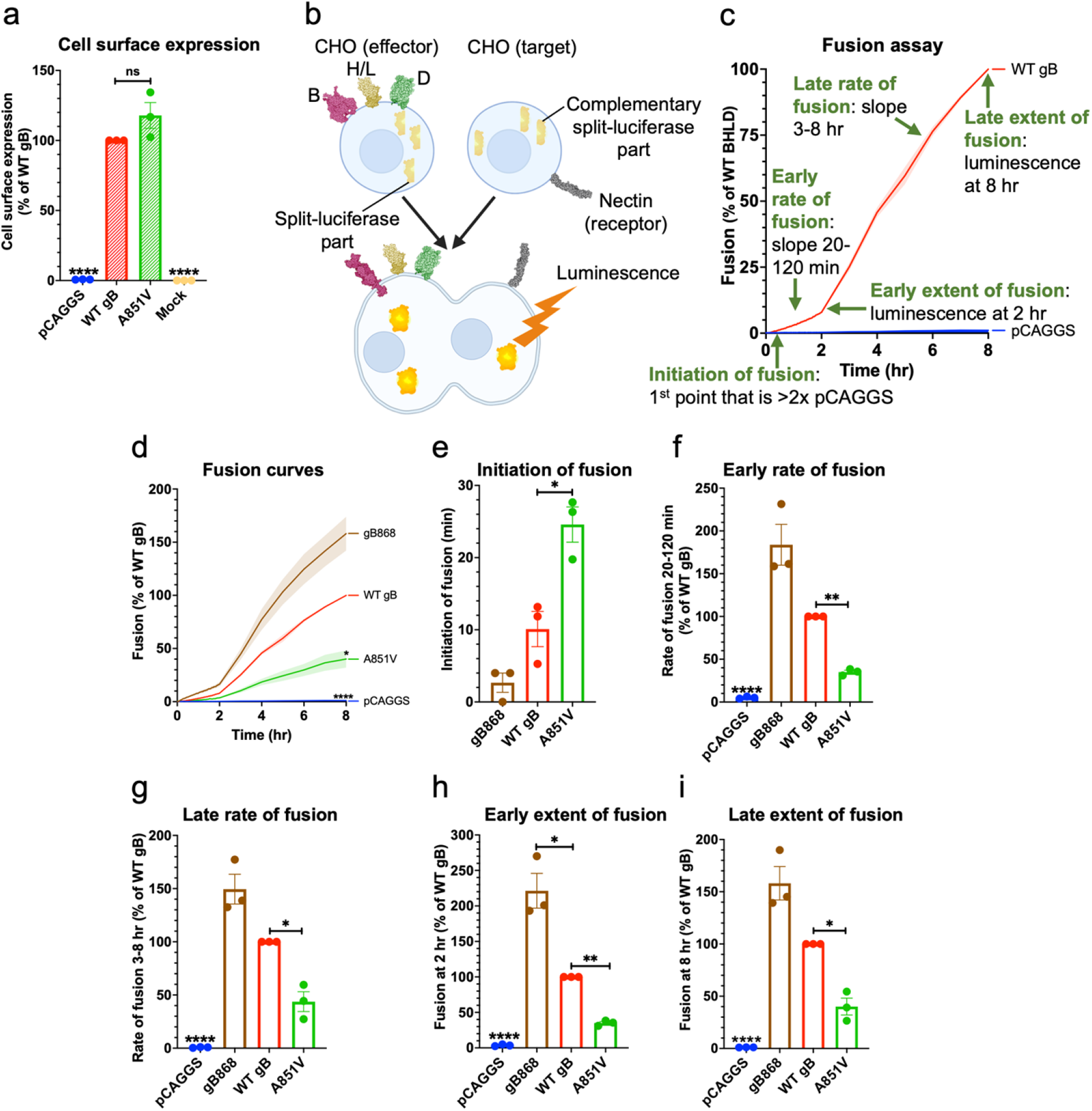
The gB_CTD_ mutant A851V reduced the rate and extent of fusion. **a)** A851V cell surface expression was not statistically different from WT gB. R68 primary antibody. Columns show mean. Error bars are SEM. **b)** Split-luciferase cell-cell fusion assay experimental setup. Cells expressing HSV-1 glycoproteins (2C36 [28], 3M1C [27], 6Z9M [2], and 5V2S [5]) fuse with cells expressing an HSV-1 receptor (3U83 [29]). Reconstitution of luciferase reports on fusion. Created with BioRender.com. **c)** Fusion of cells transfected with WT HSV-1 gB, gH, gL, gD compared to pCAGGS. The initiation of fusion is defined as the first reading at which luminescence is greater than twice that of the pCAGGS negative control. Early and late rates of fusion are the slopes of the fusion curves between 20-120 minutes and 3-8 hours post addition of target cells to effector cells, respectively. Early and late extent of fusion is defined as luminescence at 2 and 8 hours post addition of target cells to effector cells, respectively. This represents the total amount of fusion that has occurred over that time. **d)** A851V had decreased fusion compared to WT gB by the split-luciferase fusion assay. *: p < 0.05 at 8 hr. gB868 was used as a hyperfusogenic positive control [47]. Curve indicates mean values. Shaded area represents SEM. **e-i)** A851V increased the time to initiation of fusion and decreased the early and late rates and extents of fusion. Columns show mean. Error bars are SEM. *: p < 0.05, **: p < 0.01, ****: p <0.0001 in all panels. Data in all panels are from three independent experiments.

To test the hypothesis that the gB A851V mutant was hypofusogenic, we tested its fusogenicity directly. The fusogenic properties of viral fusogens are typically characterized by monitoring cell-cell fusion of uninfected cells expressing viral glycoproteins and host receptors. Here, we used a split-luciferase cell-cell fusion assay reported previously [46], in which the effector and the target cells are transfected with the complementary parts of *Renilla* luciferase. Upon fusion of effector cells with the target cells, functional luciferase forms, and the resulting luminescence is used to quantify fusion (**Fig. 2b**). This assay can measure not only early and late extent of fusion, but also early and late rate of fusion, and fusion initiation **(Fig. 2c)**.

We found that A851V mutation reduced not only the extent of fusion, but also early and late fusion rates while delaying the initiation of fusion **(Fig. 2d–i)**. The known hyperfusogenic truncation mutant gB868 was used as a control and had increased extent and rate of fusion as well as earlier fusion initiation **(Fig. 2d–i)**. The A851V fusion defect manifested within minutes and was sustained over the entire time course, reaching only ~40% of WT gB fusion by 8 hours. The fusion defect at both early and late steps in fusion suggests that A851V mutation impairs an early, rate-limiting step of fusion. Therefore, A851V is a true hypofusogenic mutation – one that impairs fusogenicity rather than folding – the first of its kind reported within the HSV-1 gB_CTD_.

### Mutational analysis of the surface pocket containing A851

A851 is located at the bottom of a surface-exposed pocket that contains another residue, T814 (**Fig. 3a–b**). The outer rim of the pocket is formed by residues L817, K807, N804, R858, A855, and L852 (**Fig. 3a**). The pocket is located at the junction of neighboring protomers such that residues N804, K807, T814, and L817 belong to one protomer, and residues A851, L852, A855 and R858 belong to the neighboring protomer (**Fig. 3a**). Thus, the gB_CTD_ trimer contains three symmetry-related pockets.

**Figure 3.**
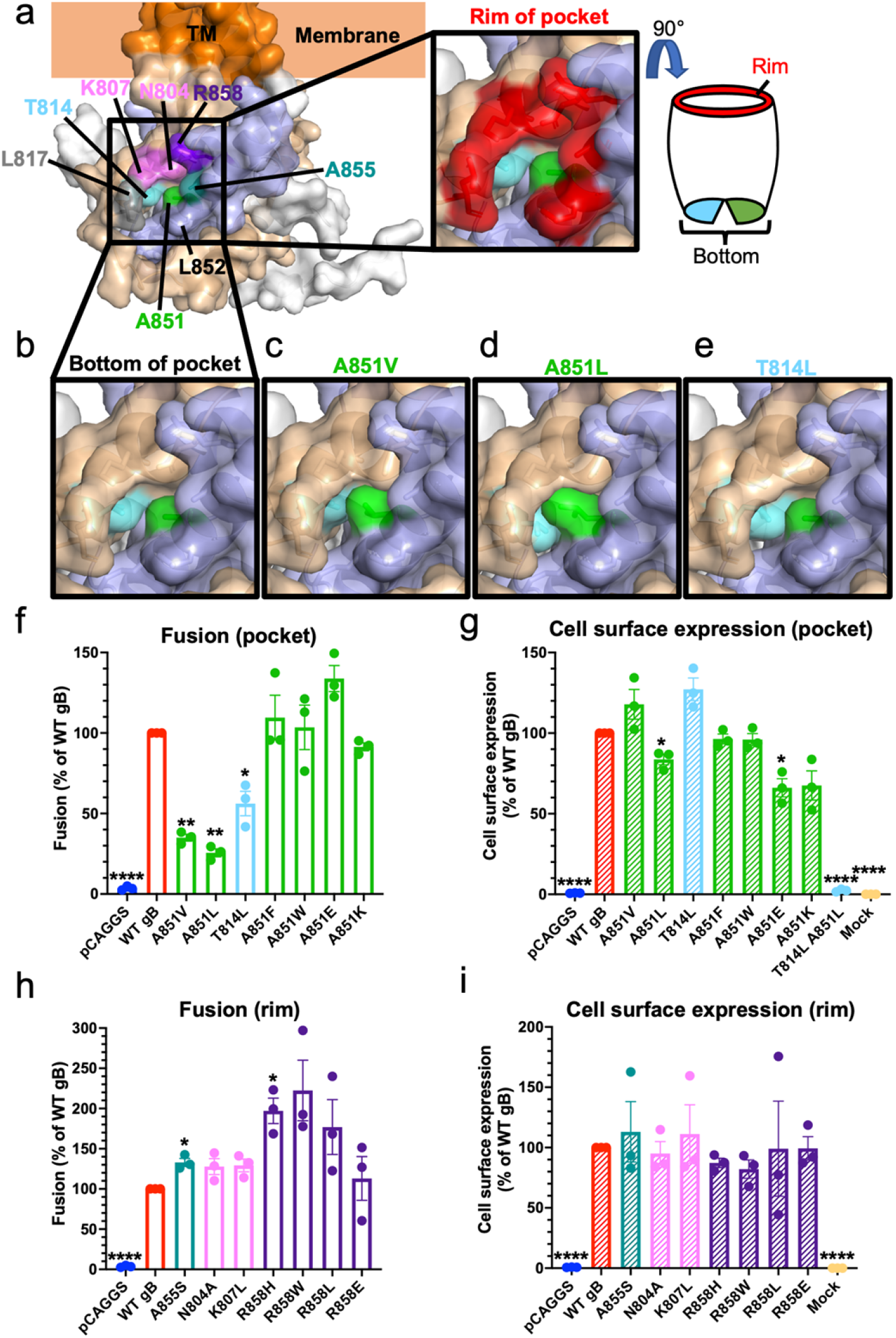
Fusogenicity and structural effects of mutants in the newly identified gB_CTD_ pocket and rim. **a)** gB_CTD_ crystal structure and the structure of the pocket on the gB_CTD_. The pocket is formed at the junction of opposing gB_CTD_ protomers, which are colored in wheat and light blue, with the third protomer in white. The residues that form the pocket on the gB_CTD_ are highlighted in colors (left panel). The residues of the outer rim of the pocket are indicated in red (right panel). **b)** The bottom of the pocket is made up of T814 and A851, in sky blue and green, respectively. They do not contact each other, leaving a space between them at the bottom of the pocket. **c-e)** Pocket mutations of T814 and A851 are hypofusogenic and were modeled in PyMOL. All three hypofusogenic mutations of T814 and A851 were predicted to fill the gB_CTD_ pocket as well as the space between T814 and A851 at the bottom of the pocket. **f)** Fusogenicity of T814 and A851 pocket mutants at 2 hr. A851V data is the same as shown in Fig. 2. Fusion trends were the same at 8 hr. **g)** Cell surface expression of pocket mutants by flow cytometry. A851V data is the same as shown in Fig. 2. **h)** Fusogenicity of mutations of the pocket rim at 2 hr. Fusion trends were the same at 8 hr. **i)** Cell surface expression of rim mutants. Columns show mean and error bars are SEM in all panels. *: p < 0.05, **: p < 0.01, ****: p < 0.0001 in all panels. Data in all panels are from three independent experiments.

We first investigated the functional significance of the pocket. A larger hydrophobic side chain of valine in the A851V mutant would reduce the size of the pocket (**Fig. 3c**). To further probe the role of A851, we mutated it to a leucine, which has a larger hydrophobic side chain than valine and would be expected to reduce the pocket size further (**Fig. 3d**). The other pocket residue, T814, was also mutated to a leucine, keeping up with the strategy of introducing a larger hydrophobic side chain **(Fig. 3e)**. Just as A851V, both A851L and T814L mutations were hypofusogenic (35%, 26%, and 56% of WT gB fusion at 2 hrs, respectively) (**Fig. 3f**). T814L had a WT-level of cell surface expression whereas the cell surface expression of the A851L mutant was slightly reduced (**Fig. 3g**).

To understand the structural basis of the observed fusion phenotypes, we examined the effect of mutations on the local structure of the gB_CTD_. At the bottom of the pocket, T814 and A851 do not directly contact one another (**Fig. 3b**). A851V, A851L, or T814L mutations are all predicted to fill the gB_CTD_ pocket **(Fig. 3c–e),** yet none are expected to destabilize the gB_CTD_ trimer. Given that all three mutations reduce fusion, the pocket appears important for fusion.

To further test this hypothesis, we designed more drastic gB pocket mutations A851F, A851W, and the double mutant T814L/A851L, to occlude the pocket fully, expecting them to decrease fusion to an even greater extent. To investigate the effect of charged residues at this position, we also designed A851E and A851K mutations. The T814L/A851L double mutant was not expressed on the cell surface (**Fig. 3g**), possibly, due to protein misfolding caused by a steric clash. A851F and A851W mutants were expressed on the cell surface at WT levels whereas A851E and A851K had slightly reduced cell surface levels **(Fig. 3g)**. Surprisingly, A851F, A851W, and A851K had no effect on fusion whereas A851E increased fusion, albeit not to a statistically significant extent (**Fig. 3f**). This was unexpected considering that all substitutions were predicted to completely fill the pocket.

Further structural modeling revealed that A851F, A851W, and A851K mutations may cause steric clashes with the surrounding residues, which could destabilize the gB_CTD_ trimer. gB_CTD_ trimer destabilization correlates with a hyperfusogenic phenotype [5]. Therefore, we hypothesized that while filling of the pocket would be expected to reduce fusion, this effect would be counteracted by the hyperfusogenic effect of trimer destabilization, yielding the observed WT fusion levels for each mutant. The charges introduced in A851E and A851K could also be causing a similar counter-effect. Collectively, our findings show that the newly identified surface pocket containing residues A851 and T814 is important for fusion (**Fig. 3a**).

### Mutational analysis of the pocket rim

We next mutated residues lining the pocket rim (**Fig. 3a**). Mutations were designed to alter side chain polarity, charge, or size (N804A, K807L, A855S, R858W, R858L, R858E) [48] while avoiding large structural changes that could destabilize the gB_CTD_ trimer because such mutations would be expected to have a hyperfusogenic phenotype [5]. Mutations were first modeled in PyMOL [49] and analyzed for significant changes to the surrounding gB_CTD_ structure. Previously reported hyperfusogenic mutations L817H [33] and L817P [34] were not tested here. L852 was not mutated because its mutation L852A was reported to abrogate cell surface expression [32].

Mutations of the rim residues were either hyperfusogenic or had no effect on fusion (**Fig. 3h**), and all had similar cell-surface expression as WT gB (**Fig. 3i)**. A855S was moderately hyperfusogenic, which is consistent with the hyperfusogenic phenotype of A855V reported by others [33, 38]. N804A and K807L were also moderately hyperfusogenic but did not reach a statistically significant level. R858H (also tested in [35, 40, 42, 47]), R858W, and R858L were markedly hyperfusogenic, judging by the mean values, even though increased fusion levels of R858W and R858L were not statistically significant due to large SEM values. This is consistent with the hyperfusogenic phenotype of R858C reported by others [31]. Finally, R858E had no effect on fusion. According to structural modeling, most mutations are predicted to expose the pocket opening (L817P, N804A, K807L) or the protomeric interface (R858C, R858H, R858L, R858E) whereas some mutations do not (A855S, A855V, and R858W). R858 crosses the fault line (protomeric interface) between adjacent protomers but lies in the upper rim of the pocket and does not cover the pocket (**Fig. 3a**). Therefore, many of its mutations are predicted to expose the protomeric interface without altering the exposure of the pocket opening directly. Many mutations also neutralize a positive charge in the upper portion of the rim (K807L, R858C, R858H, R858W, and R858L). While the predicted structural effects are diverse, the mutations in the rim of the gB_CTD_ pocket are overwhelmingly hyperfusogenic, with the exception of R858E, and the observed hyperfusogenic phenotypes correlate with a greater exposure of the pocket or the protomeric interface and neutralization of a positive charge. This suggests a regulatory role for the rim in the fusogenicity of gB.

### gH V831 is the most important gH_CT_ residue for fusion

Mutations that reduce the size of the A851/T814 pocket in gB_CTD_ without destabilizing the trimer – A851V, A851L, and T814L – reduce fusion. Therefore, the size of the pocket is important for fusion, and we hypothesized that it could function as a binding site. Previously, we showed that gH_CT_ has an activating role in fusion because its truncations progressively reduce fusion [42], and proposed that gH may activate gB by using gH_CT_ to disrupt the inhibitory gB_CTD_ trimer in a wedge-like manner [5]. Therefore, we hypothesized that the gB_CTD_ pocket was the binding site for the gH_CT_.

To identify the gH_CT_ residues that could interact with the gB_CTD_ pocket, we first narrowed down residues within the 14-residue gH_CT_ necessary for fusion. In our previous work, we showed that the gH832 truncation mutant lacking 6 residues of the gH_CT_ (**Fig. 4a–b**) had WT-level fusion whereas the gH829 mutant lacking 9 residues had a significantly reduced fusion [42]. To narrow down residues most important for fusion, we tested serial truncations of gH829, gH830, gH831, and gH832 **(Fig. 4b)**. gH832 was slightly hyperfusogenic (**Fig. 4c**) whereas gH831, gH830, and gH829 were hypofusogenic, with fusion extent proportional to the length of the remaining gH_CT_ **(Fig. 4c–d).** Cell surface expression of the gH truncation mutants was similar to the WT gH expression (**Fig. 4h**).

**Figure 4.**
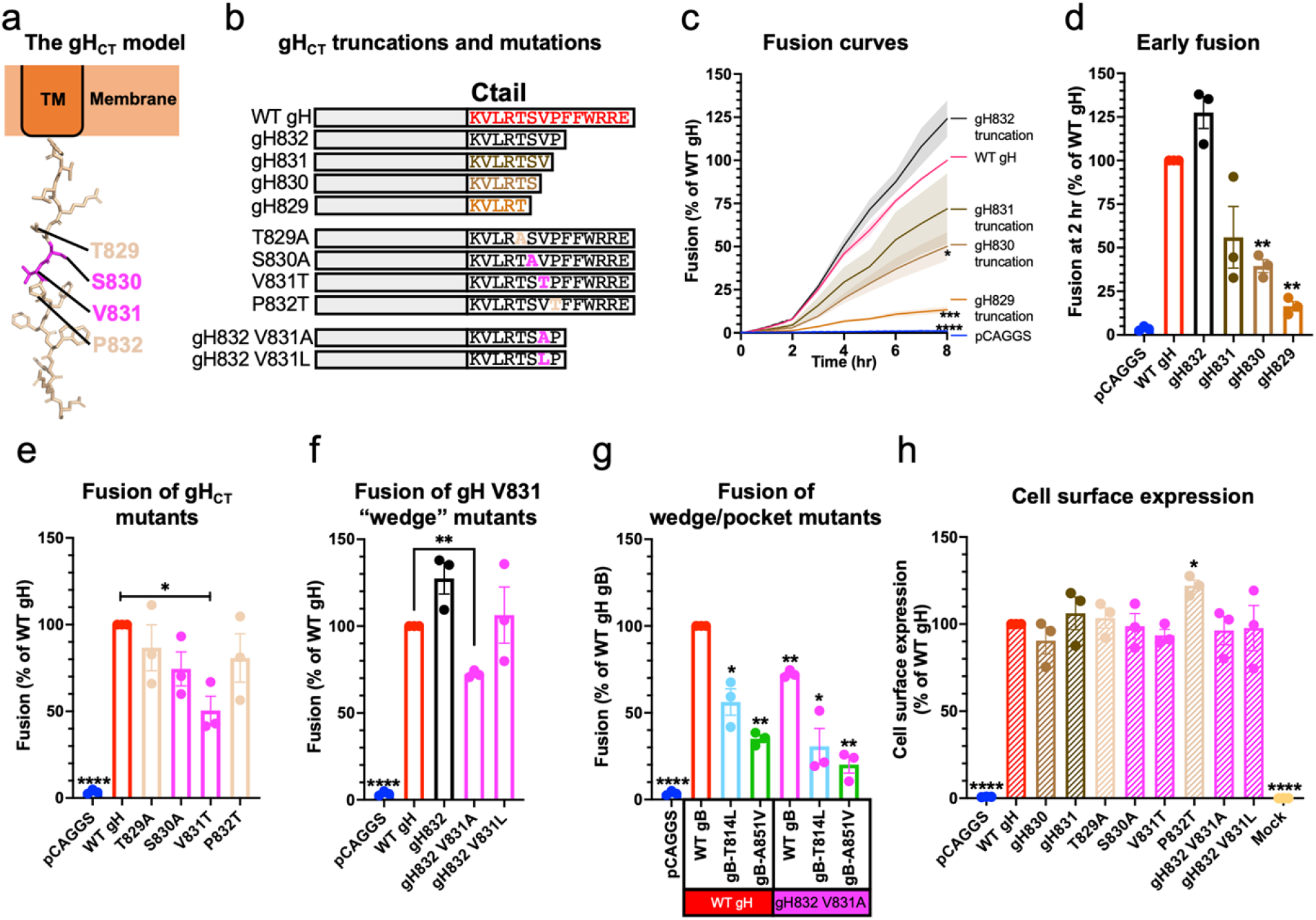
gH V831 is the most important gH_CT_ residue for fusion. **a)** Structural modeling of the gH_CT_. **b)** Summary of gH_CT_ truncations and mutations tested to determine which gH_CT_ residues are the most important for fusion and probe their mechanism of action. **c)** Kinetics of fusion of gH_CT_ truncations over 8 hr. Statistical significance shown is based on comparisons to WT gH fusion at 8 hr. Curves are the mean. Shaded area is SEM. **d)** Fusion of gH_CT_ truncations at 2 hr. **e)** Fusion of mutations of gH829-832 residues at 2 hr to probe the function of the residues. Fusion trends were the same at 8 hr. **f)** Fusion of V831 mutations designed to make the putative gH wedge smaller (V831A gH832) or larger (V831L gH832), at 2 hr. Fusion trends were the same at 8 hr. **g)** Fusion of mutations creating a smaller wedge (V831A gH832) combined with mutations creating smaller pockets (gB T814L, gB A851V), at 2 hr. Fusion trends were the same at 8 hr. **h)** Cell surface expression of the gH_CT_ truncations and mutations. Expression of gH829 and gH832 was determined previously to be the same as WT gH expression [42]. LP11 primary antibody. *: p < 0.05, **: p < 0.01, ***: p < 0.001, ****: p < 0.0001 in all panels. Bars indicate the mean and error bars are SEM. Statistical comparisons are to WT gH and gB. All panels represent averages of three independent experiments.

To further probe the functional importance of residues T829, S830, V831, and P832, we reversed their polarity [48] by either making them more hydrophobic (T829A, S830A, and P832T) or more hydrophilic (V831T), in the context of the full-length gH (**Fig. 4b**). Both S830A and V831T were hypofusogenic, with V831T having the greater, statistically significant fusion defect (**Fig. 4e**). There were no significant differences in cell surface expression relative to WT gH, except for P832T, which had slightly increased expression (**Fig. 4h**). Collectively, these data implicated V831 as the most important gH_CT_ residue for fusion.

The gH_CT_ is predicted to be unstructured, and when it was modelled as an extended polypeptide, gH V831 ended up approximately the same distance from the membrane as the gB_CTD_ binding pocket (**Fig. 5a**). This means that when gH and gB come into proximity, gH V831 could, in principle, interact with the gB_CTD_ binding pocket (**Fig. 5a**). Therefore, we propose that V831 is the gH_CT_ “wedge” and that it binds the gB_CTD_ pocket containing T814 and A851 (**Fig. 5a**).

**Figure 5.**
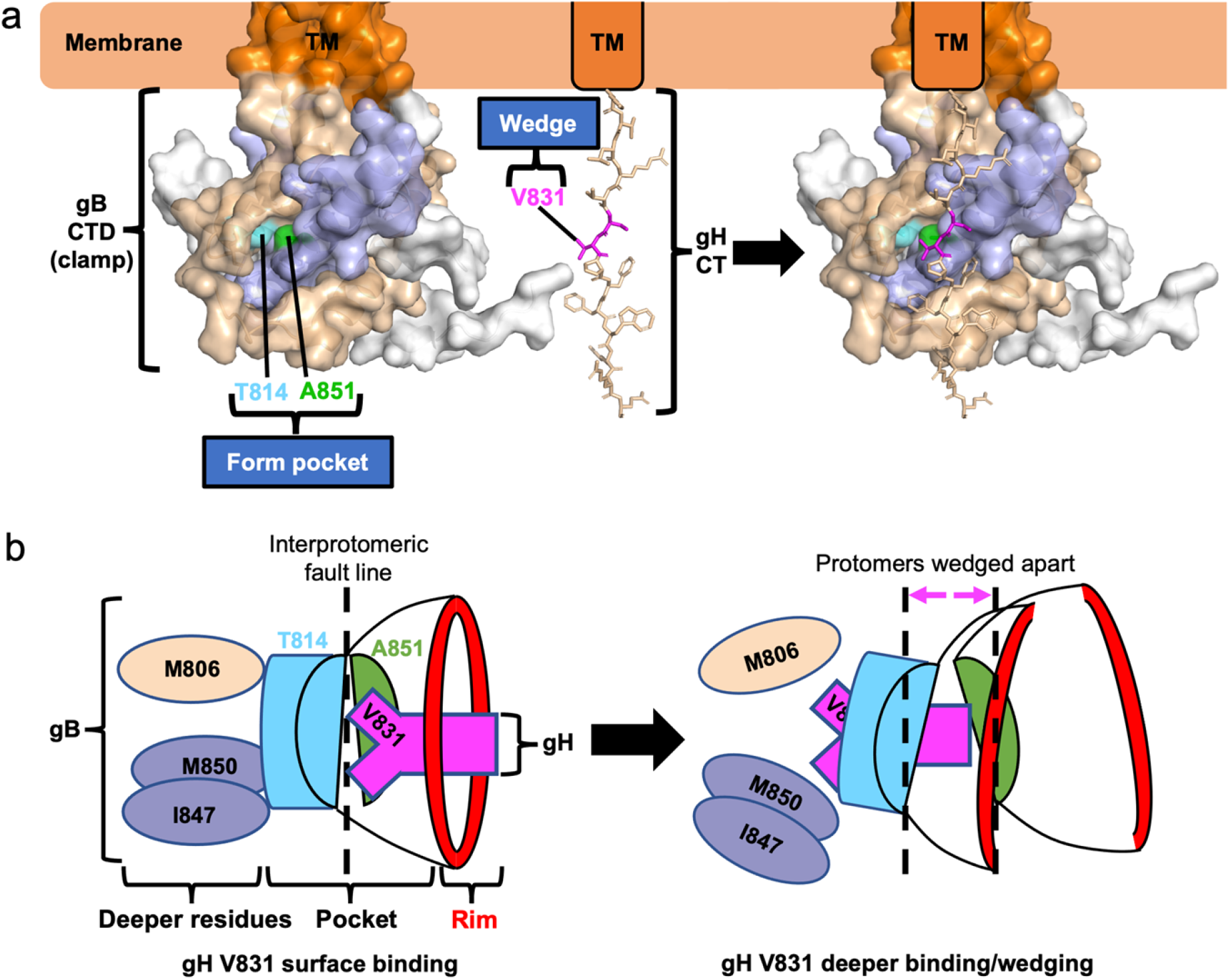
A model for gB_CTD_/gH_CT_ interactions and fusion triggering. **a)** gB_CTD_ residues T814 (sky blue) and A851 (green) form a gH-binding pocket on the surface of the gB_CTD_ trimer. gH V831 (magenta) acts as a wedge. Modeling shows that the gH V831 wedge and the gB_CTD_ pocket are equidistant from the membrane. The V831 wedge binds between gB T814 and A851 in the gB pocket and pushes the gB_CTD_ protomers (wheat and light blue) apart to destabilize gB and trigger fusogenic refolding of gB into the postfusion conformation. **b)** gH V831 (magenta) acts as a wedge that initially binds between gB_CTD_ residues T814 (sky blue) and A851 (green) in the pocket on the surface of the gB_CTD_ trimer. gH V831 then binds to deeper hydrophobic residues of the gB_CTD_ (colored by protomer), forming favorable hydrophobic interactions. This causes the protomers of gB to be pushed apart as gH V831 enters deeper into the gB_CTD_, destabilizing the gB_CTD_ clamp and triggering gB to refold and cause fusion.

We next investigated the side chain requirement at the gH residue 831. A hydrophobic side chain appears to be required at this location because threonine could not effectively substitute for valine despite a comparable side chain size (**Fig. 4b and e**). We then asked whether a smaller (alanine) or a larger (leucine) hydrophobic side chain could support efficient fusion.

Previously, we showed that V831A mutation was hypofusogenic [50]. Thus, a smaller hydrophobic side chain of alanine could not substitute for a valine. Interestingly, the hypofusogenic phenotype of V831A was observed only in the context of a truncated gH832 but not full-length gH [50], suggesting that residues 833-838 somehow compensated for the fusion defect of the V831A mutation. To test the effect of a larger hydrophobic side chain at position 831, we generated the V831L mutation in the context of the gH832 truncation (gH832 V831L) to prevent potential compensation by residues 833-838 (**Fig. 4b**). gH832 V831L had WT fusion level (**Fig. 4f**) whereas gH832 V831A was hypofusogenic, in accordance with our previous finding [50]. Neither gH832 V831A nor gH832 V831L mutations had any effect on cell surface expression (**Fig. 4h**). We conclude that valine is minimally required at position 831 to maintain WT levels of fusion.

We then tested the possibility that, in the context of a smaller gB_CTD_ pocket (gB A851V or T814L), a smaller gH_CT_ wedge (gH832 V831A) could support fusion by being able to fit into a smaller pocket. We found that the small pocket/small wedge combinations did not restore fusion to WT levels but, instead, reduced fusion in an additive manner (**Fig. 4g**). Therefore, we hypothesize that both the gB_CTD_ pocket and the gH_CT_ wedge have to be of a certain size to function efficiently.

## DISCUSSION

### Identification of a new functional pocket in the gB_CTD_

Using mutational analysis, we have identified a new functional region of the gB_CTD_ composed of a surface-exposed pocket and its rim (**Fig. 3a**). The bottom of the pocket is formed by residues T814 and A851 and the rim is formed by residues N804, K807, L817, L852, A855, and R858. The pocket is located at the junction of two gB protomers, and the pocket and the rim residues are evenly distributed between the neighboring protomers, with N804, K807, T814, and L817 located in one protomer and A851, L852, A855, and R858 in the neighboring protomer. Thus, there are three such pockets within the gB_CTD_ trimer.

Mutations of the gB_CTD_ pocket and rim had opposite effects on fusion. Pocket mutations A851V, A851L, and T814L, which partially filled the pocket without otherwise perturbing the surrounding gB_CTD_ structure, significantly decreased fusion, i.e., were hypofusogenic. These three mutations are unusual because, to the best of our knowledge, they are the first hypofusogenic HSV-1 gB_CTD_ mutations that perturbed protein function rather than caused protein misfolding. The hyperfusogenic rim mutations are also unusual because they differed from most other hyperfusogenic mutations in both their location and the presumed mechanism of action. Most hyperfusogenic mutations in the gB_CTD_ [31–43] map to either the protomeric interfaces or the membrane-binding regions and are thus predicted to destabilize the membrane-dependent gB_CTD_ trimer [5]. In contrast, the hyperfusogenic rim mutations, which include mutations of residues L817, A855, and R858 reported here and elsewhere [31, 33–35, 38, 40, 42, 47], are located on the surface and would not be predicted to disrupt the gB_CTD_ trimer. The gB_CTD_ pocket is also located too far from the membrane to participate in membrane interactions. Therefore, the rim mutations increase fusion by a different mechanism, potentially, by exposing the pocket (see below). Collectively, our mutational analysis and structural modeling uncovered a new fusogenic site within the gB_CTD_ trimer composed of two distinct functional regions, the pocket and its rim.

### The gB_CTD_ pocket is a putative binding site for the gH_CT_ wedge, residue V831

Surface pockets often function as binding sites, and mutations that fill surface pockets typically disrupt protein function by blocking binding to protein partners. For example, mutagenesis of a putative chaperone-binding pocket of the p53 protein decreased its expression, indicating that the binding pocket was important for binding the chaperone that stabilizes p53 [51]. Introducing an amino acid with a bulkier, branched side chain into a pocket on Rabies virus P protein resulted in decreased binding by STAT proteins, suggesting that the mutated pocket constitutes the STAT-binding site [52]. Furthermore, in enzymes, active sites, which bind and convert substrates, are typically found in surface pockets, and pocket-filling mutations can either block binding of the substrate to the active site [53] or obstruct its access through a tunnel or channel [54, 55].

Mutations that filled the gB_CTD_ pocket decreased fusion, suggesting that the size of the pocket is important for fusion, so we hypothesized that the gB_CTD_ pocket serves as a binding site for a protein partner and that the binding event activates fusion. Previously, we speculated that gH may activate gB by using gH_CT_ to disrupt the inhibitory gB_CTD_ trimer in a wedge-like manner [5]. Therefore, we hypothesized that the newly identified gB_CTD_ pocket was the binding site for the gH_CT_. Using mutational analysis, we identified gH_CT_ residue V831 as the single most critical residue for fusion. Our structural analysis predicted that if the 14-residue gH_CT_ adopted an extended conformation, gH V831 and the gB_CTD_ pocket would end up roughly equidistant from the membrane, putting them into ideal positions for reciprocal interaction (**Fig. 5a**). Based on these results, we propose that V831 binds in the gB_CTD_ pocket (**Fig. 5a**).

We hypothesize that to trigger fusion, the gH V831 side chain binds the gB_CTD_ pocket and inserts deeply enough to push the gB protomers apart like a wedge or a crowbar. This destabilizes the gB_CTD_ clamp, which releases its inhibitory hold on the gB ectodomain, allowing the latter to refold from the prefusion into the postfusion conformation. This hypothesis is supported by the location of the gB_CTD_ pocket right above the interprotomeric “fault line”, a prime location for pushing the protomers apart. Just as the size of the gB_CTD_ pocket is critical for fusion (with a smaller pocket decreasing fusion), so is the size of the sidechain at gH residue 831. A smaller alanine (V831A) decreased fusion whereas a larger leucine (V831L) preserved WT-level fusion. Thus, there appears to be a minimum requirement for the size of the side chain at gH residue 831 for WT-levels of fusion. An alanine would not be able to insert deeply enough between the gB protomers and would be less effective at destabilizing the gB_CTD_, which explains why V831A mutant is hypofusogenic. Interestingly, V831T mutant was also hypofusogenic. The side chain of threonine is similar in size to valine yet is more hydrophilic, which suggests that in addition to size, hydrophobicity of the gH residue 831 is important for fusion.

Implicit in our insertion model is the assumption that the gH V831 residue first interacts with T814 and A851 as it inserts into the pocket and then may interact with nearby residues located within the gB_CTD_ core as it wedges in deeper to push the protomers apart (**Fig. 5b**). The residues that lie underneath the gB_CTD_ pocket, M806, I847, and M850, are hydrophobic, so a hydrophobic residue would be a more effective wedge because it could form more favorable interactions with these residues (**Fig. 5b**). This helps explain why a slightly larger and hydrophobic leucine (V831L) is an efficient substitute for the native valine whereas a similarly sized yet hydrophilic threonine (V831T) is not. The observation that the residues underneath the gB_CTD_ pocket are hydrophobic further supports our model that the pocket is the binding site for the gH_CT_ wedge, residue V831.

### Role of the rim of the gB pocket in fusion

In contrast to the hypofusogenic mutations of the gB_CTD_ pocket, nearly all mutations of the pocket rim were hyperfusogenic. This included both mutations made in this work (R858W, R858L, R858E, A855S), and those reported previously (R858H [35, 40], R858C [31], L817H [33], L817P [34], A855V [33, 38]). Being located on the surface, none of these mutations would be predicted to disrupt the gB_CTD_ trimer, in contrast to the majority of the known hyperfusogenic mutations [5]. Therefore, the pocket rim mutations enhance fusion by a different mechanism.

The rim is the entryway into the gB_CTD_ pocket, and many of the hyperfusogenic rim mutations (N804A, K807L, R858C, R858H, R858W, and R858L) appear to expose the entryway into the pocket to some extent. This may increase fusion by facilitating access of the gH_CT_ wedge to the gB_CTD_ pocket. Many rim mutations neutralize the positive charge in the upper portion of the pocket (K807L, R858C, R858H, R858W, and R858L), which also correlates with hyperfusogenicity. R858E was the only rim mutation that was fusion-neutral. While it is predicted to expose the entryway into the pocket, R858E mutation replaces the positive charge with a negative one. Thus, neutralization of a positive charge in the upper portion of the pocket rim could facilitate access of the gH_CT_ wedge to the pocket. Congruent with this idea is that the gH_CT_ is mostly uncharged, notably, residue V831.

The A855S and A855V mutations do not change the charge or the access to the pocket. Nonetheless, we hypothesize that by analogy with the other rim mutations listed above, A855S and A855V mutations somehow facilitate gH_CT_ interactions with the gB_CTD_ pocket. The hyperfusogenic phenotype of L817P and L817H mutants is difficult to explain due to insufficient structural data; L817 is the last resolved residue before a disordered loop and its conformation is likely dynamic and difficult to predict.

### More drastic gB A851 pocket mutations may offset the pocket-filling effect with trimer destabilization

Having determined that the gB_CTD_ pocket-reducing mutations A851V, A851L, and T814L all reduced fusion, we had anticipated that more drastic mutations would reduce fusion further. Therefore, we designed mutations A851F and A851W to completely fill the gB_CTD_ pocket and A851E and A851K to both fill the pocket and introduce charge. However, A851F, A851W, and A851K were fusion-neutral, and A851E was slightly hyperfusogenic. Structural analysis suggested that due to their large side chains, both phenylalanine and tryptophan would clash with nearby residues causing significant steric strain. Since the gB_CTD_ pocket spans neighboring protomers, and mutations predicted to disrupt the gB_CTD_ trimer are hyperfusogenic [5], we hypothesize that in addition to filling the gB_CTD_ pocket, A851F or A851W also destabilize the gB_CTD_ trimer. As a result, the hypofusogenic phenotype expected of the pocket-filling mutation is counterbalanced by the hyperfusogenic effect of gB_CTD_ destabilization, resulting in the observed WT-level fusion phenotype. gB mutations A851E and A851K would likewise be predicted to destabilize the gB_CTD_ trimer due to steric strain and, additionally, due to introduction of a charge.

### Open questions

According to our model of gB triggering, residue V831 within the gH_CT_ acts as a wedge that binds in the gB_CTD_ pocket and pushes the gB protomers apart. This destabilizes the gB_CTD_ trimer, relieving its inhibitory hold on the ectodomain and allowing fusogenic refolding. However, a few unanswered questions remain. First, we do not yet have direct evidence of this gB/gH interaction. Second, it is unknown how the signal from the destabilized gB_CTD_ trimer is transmitted to the gB ectodomain on the other side of the membrane. A recent study suggested that a conserved regulatory helix in the gB ectodomain proximal to the membrane may be involved in the transduction of the triggering signal from the cytoplasmic domain to the ectodomain [56]. Further studies are needed to decipher how gH/gL interacts with and activates gB and how the gB structure changes in response to this interaction.

## MATERIALS AND METHODS

### Cells and plasmids

CHO cells were received as a gift from J. M. Coffin and grown in Ham’s F-12 medium supplemented with 10% fetal bovine serum (FBS), 100 IU penicillin, and 100 μg/ml streptomycin at 37° C in the presence of 5% CO2, except as noted otherwise. Plasmids pPEP98, pPEP99, pPEP100, and pPEP101 contain the full-length HSV-1 (strain KOS) gB, gD, gH, and gL genes, respectively, in a pCAGGS vector and were gifts from P. G. Spear [57]. Plasmids RLuc1-7 and RLuc8-11 (carrying the *Renilla* split luciferase genes) and pBG38 (carrying the nectin-1 gene) were gifts from G. H. Cohen and R. J. Eisenberg [46, 58]. Plasmids pJLS11 (gB868) [47], pJLS8 (gB R858H) [47], pJLS15 (gH832) [50], pJLS16 (gH832 V831A) [50], and pHR26 (gH829) [42] were generated previously in our lab.

### gB_CTD_ mutagenesis

Point mutations in the cytoplasmic domain of the full-length gB gene were generated in pPEP98 background by using PCR and Gibson assembly [59]. pPEP98 was cut at PmlI and MfeI sites to generate the backbone for the assembly. Two DNA fragments were created by PCR for the mutants using the primers listed in Table 1. The digested backbone and two DNA inserts generated by PCR were assembled using the Gibson Assembly Master Mix from New England Biolabs. The T814L/A851L double mutant was generated using the same strategy, by making a plasmid with the T814L mutation first and then repeating the steps to introduce the A851L mutation. The A851E mutant was generated by QuikChange PCR [60] using the primers listed in Table 1 followed by ligation with T4 ligase. The resulting plasmids were pZP6 (A851V), pZP22 (N804A), pZP23 (K807L), pZP7 (T814L), pZP21 (A851L), pZP25 (A851K), pZP25 (A855S), pZP60 (R858W) pZP24 (R858L), pZP27 (R858E), pZP57 (A851F), pZP58 (A851W), pZP61 (T814L/A851L), pZP2 (A851E).

**Table 1.**
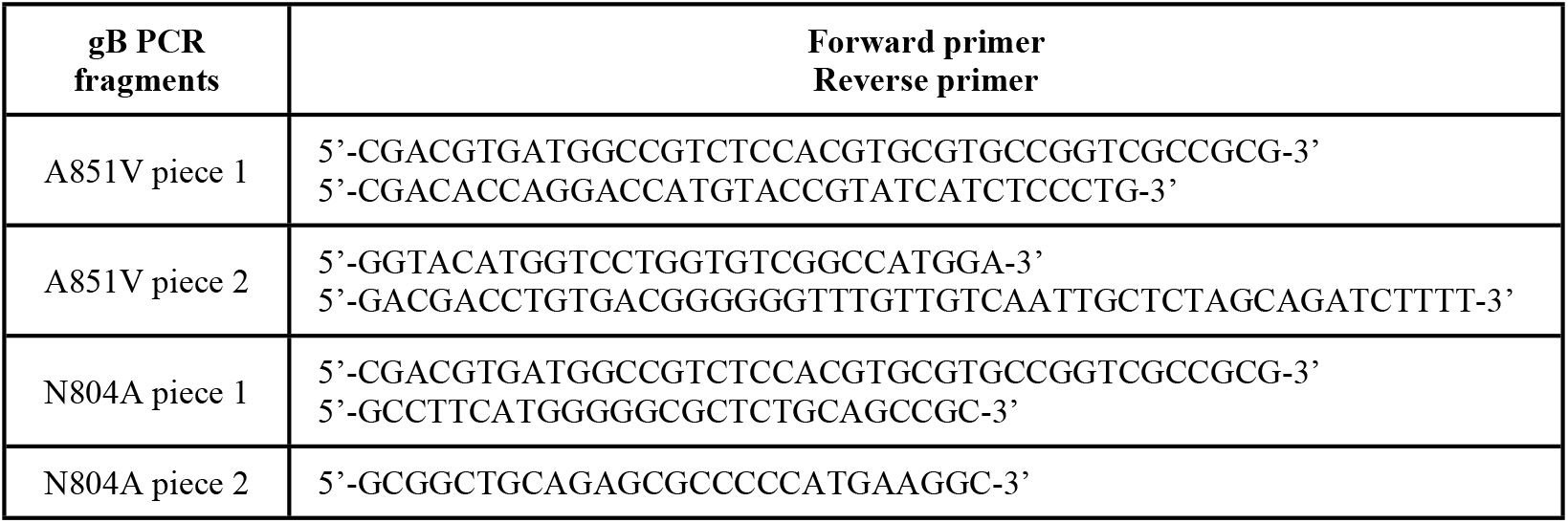
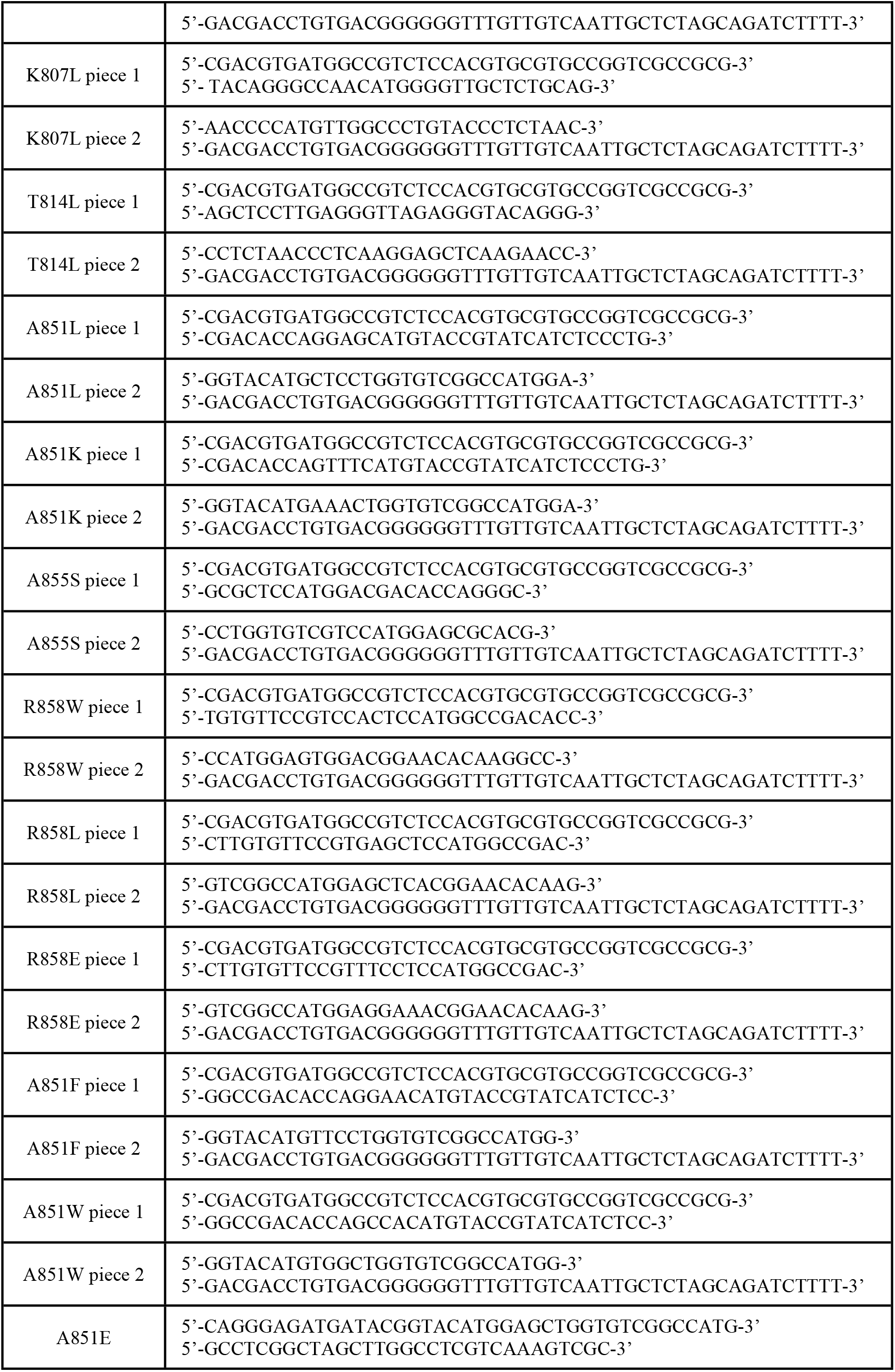
Primers used for gB_CTD_ mutagenesis.

### gH_CT_ mutagenesis

Point mutations in and truncations of the gH_CT_ were generated in the pPEP100 background by using PCR and Gibson assembly [59]. pPEP100 was cut at MfeI and XhoI sites to generate the backbone for the assembly. Two DNA fragments were created by PCR for the mutants using the primers listed in Table 2. The digested backbone and two DNA inserts generated by PCR were assembled using the Gibson Assembly Master Mix. gH832 V831L was constructed by digesting pJLS15 with MfeI and XhoI to obtain the backbone for the assembly, PCR of the insertion fragment using the primers listed in Table 2, and assembly using the Gibson Assembly Master Mix. The resulting plasmids were pZP32 (gH830), pZP33 (gH831), pZP28 (T829A), pZP29 (S830A), pZP30 (V831T), pZP31 (P832T), pZP62 (gH832 V831L).

**Table 2.**
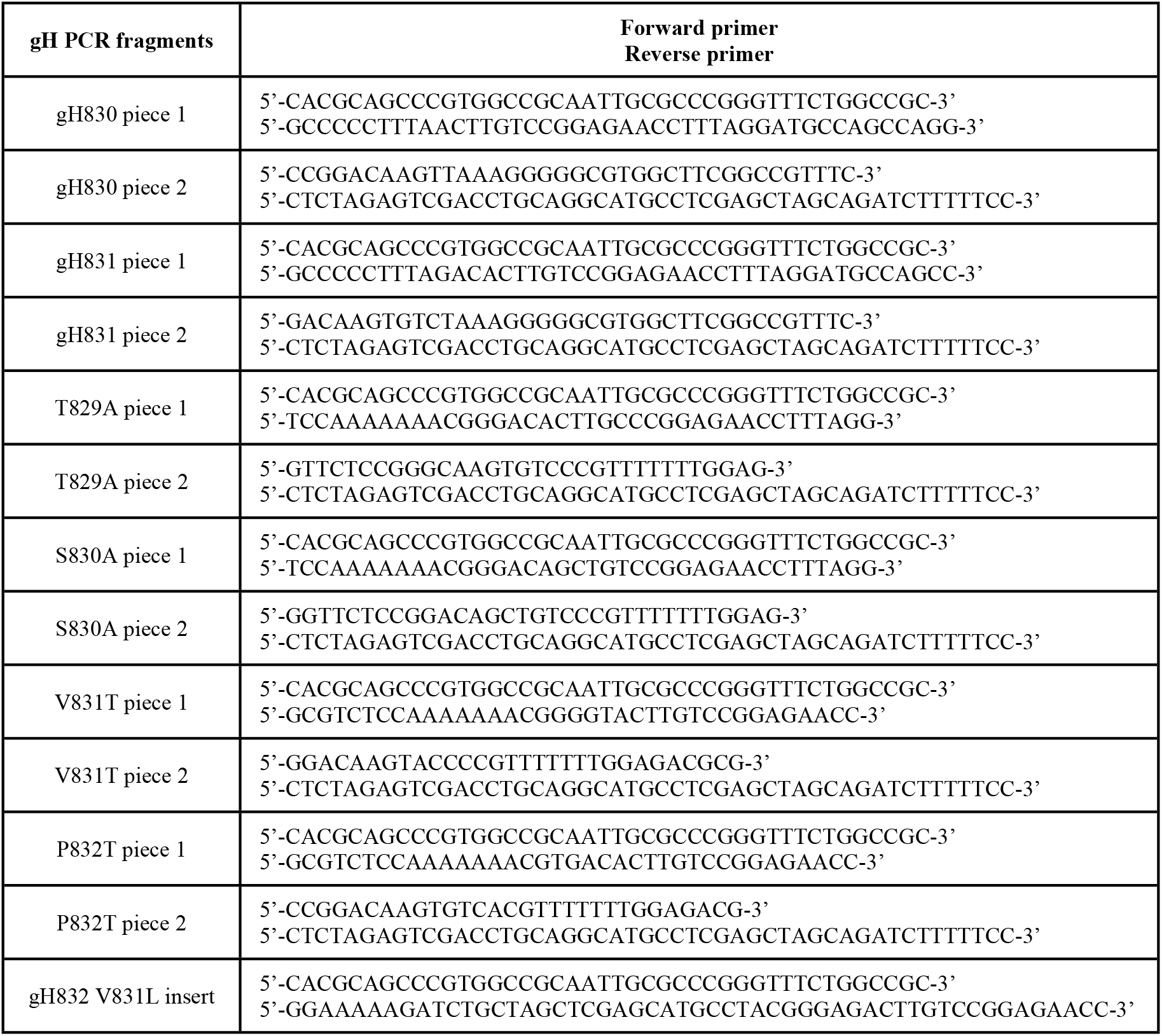
Primers used for gH_CT_ mutagenesis.

### Cell-cell fusion assay

Cell-cell fusion of gB and gH mutants was measured using a split-luciferase assay [46]. Chinese hamster ovary (CHO) cells [61] were seeded into 96-well plates at 50,000 cells per well, three wells per condition, for effector cells and 6-well plates at 200,000 cells per well for target cells. The next day, effector cells were transfected per well with 125 ng gB (pPEP98 or gB mutant) and 41.7 ng each of split luciferase (RLuc1-7), gH (pPEP100 or gH mutant), gL (pPEP101), and gD (pPEP99) using 0.58 μl JetPrime (Polyplus, Illkirch-Graffenstaden, France) in 10 μl JetPrime buffer. For the pCAGGS negative control condition, 250 ng of pCAGGS empty vector was transfected in place of the gB, gH, gL and gD plasmids. CHO cells lack HSV-1 receptors, so no fusion can occur until receptor-bearing target cells are introduced [62]. Each well of target cells was transfected with 1 μg of the complementary part of the split luciferase (RLuc8-11) and 1 μg of the HSV-1 receptor nectin-1 (pBG38) with 4 μl of JetPrime in 200 μl of JetPrime buffer. On day 3, the tissue culture media was removed from the 96-well plate wells and replaced with 40 μl per well of fusion medium (Ham’s F12 with 10% FBS, Penicillin/Streptomycin, 50 mM HEPES), with 1:500 Enduren luciferase substrate (Promega, Madison, WI) added. The Enduren concentration becomes 1:1000 once the target cells are added. Cells were incubated for 1 hr at 37° C. Meanwhile, target cells were detached by incubating with 1 ml per well of Versene (Fisher Scientific, Waltham, MA). Target cells were collected, spun down, and resuspended in 500 μl of fusion medium per well. 40 μl of target cells were added to each well of effector cells. The plate was immediately placed in a BioTek plate reader. Luminescence measurements were taken every 1-2 minutes for 2 hrs followed by measurements every hour until hour 8. Either gB868 or gB R858H was always included as a hyperfusogenic positive control to ensure that the assay was working as expected. The average hyperfusogenic positive control signal was higher than that of the WT condition in all experiments. Luminescence values were then averaged for the three wells in each condition, normalized to the WT signal at 8 hrs, and expressed as a percentage of WT. Reported values are averages of three biological replicates.

### Flow cytometry

Cell surface expression of gB and gH mutants was measured using flow cytometry. CHO cells were seeded at 250,000 cells per well in 6-well plates. The next day, each well was transfected with 2 μg gB (pPEP98 or gB mutant), or pCAGGS, or 1 μg each of gH (pPEP100 or gH mutant) plus 1 μg gL (pPEP101) using 4 μl of JetPrime in 200 μl JetPrime buffer. One well per experiment was left untransfected as a ‘mock’ control. On day 3, the cells were detached with 1 ml per well of Versene and collected using FACS medium (PBS with 3% FBS). Cells were washed with FACS media and incubated for 1 hr on ice with 250 μl of primary antibody (R68 for gB and pCAGGS, LP11 for gH and pCAGGS, anti-c-myc rabbit (A14, Santa Cruz Biotechnology, Dallas, TX) or mouse (9E10, Santa Cruz Biotechnology) antibody as a non-targeting negative control for the Mock condition) at a 1:500 dilution in FACS medium. Cells were washed three times and incubated for 1 hr on ice in the dark with 250 μl secondary FITC-conjugated anti-rabbit antibody (for R68 and rabbit anti-myc; MPBio, Santa Ana, CA) or Alexa Fluor 488 anti-mouse antibody (for LP11 and mouse anti-myc; Invitrogen, Waltham, MA) at a 1:250 dilution in FACS medium. Cells were washed three times and resuspended in 500 μl of FACS medium. Cell fluorescence was measured by flow cytometry. Gating of live cells was performed based on FSC and SSC using FlowJo software. gB+ and gH+ cells were gated using the pCAGGS condition as a negative control, using a cutoff of 5% gB+ or gH+ pCAGGS cells to capture the vast majority of true positives while minimizing false positives. Total cell surface expression of the transfected population was obtained by calculating the product of % gB+ or gH+ cells and the mean fluorescence intensity of the gB+ or gH+ cells. Total cell surface expression was then normalized to the WT gB or WT gH condition, expressed as a percentage. The values represent the average of three independent experiments. R68 (polyclonal anti-HSV-1 gB) was a gift from G. H. Cohen and R. J. Eisenberg. LP11 (monoclonal anti-HSV-1 gH/gL) was a gift from Helena Browne.

### Statistics

Statistical analysis was performed for each experiment on the normalized values using GraphPad PRISM 9 software. Unpaired t-test with Welch’s correction was used to compare conditions to each other as indicated.

### Structural analysis

The crystal structure of full-length HSV-1 gB, PDB 5V2S [5] was used for *in silico* structural analysis. Predicted effects of gB_CTD_ mutants on the structure were analyzed using PyMOL software ([49] Version 2.5.1 Schrödinger, LLC.). Mutations were introduced one by one, selecting the rotamer with the highest probability and lowest clashing with nearby residues. The area where the mutation was introduced was then “cleaned” in a 5-Å radius using the ‘clean’ function in PyMOL, which analyzes the selected area and shifts residues into positions that are predicted to be more favorable to minimize energy and strain. The resulting structure after cleaning is more likely to reflect the actual structure of the mutant. The “cleaned” model of the mutant was then visually compared to the WT structure. Predicted changes to salt bridges and H-bonds were also analyzed. The gH_CT_ was modelled as an unstructured peptide in PyMOL. The electrostatic surface potential of the gH_CT_ was calculated using the PyMOL ABPS Tools v. 2.1.5 plugin.

## ACKNOWLEDGEMENTS

We thank Stephen Kwok and Allen Parmelee from the Tufts Flow Cytometry Core and Adam Hilterbrand for flow cytometry training. We thank Roselyn Eisenberg and Gary Cohen (U. Pennsylvania) for the gift of the antibodies and split luciferase plasmids. We thank Doina Atanasiu (U. Pennsylvania) for advice regarding the split luciferase fusion assay. We thank John Coffin, Marta Gaglia, and Karl Munger for helpful discussions and Andrea Rebolledo Viveros for comments on the manuscript. PyMOL software was installed and maintained by SBGrid [63]. This work was funded by the grants T32GM731042 (Z.P.), T32GM731043 (Z.P.), F30AI161795 (Z.P.), and 1R01AI164698 (E.E.H.) from the National Institutes of Health, and by a Faculty Scholar grant 55108533 from Howard Hughes Medical Institute (E.E.H.).

## AUTHOR CONTRIBUTIONS

Z.P. designed the experiments, cloned the constructs, conducted the experiments, analyzed the data, performed structural analyses, generated hypotheses, generated models, and wrote the manuscript. E.K.S. conducted experiments, analyzed the data, performed structural analyses, and generated hypotheses. E.E.H. designed experiments, analyzed the data, generated hypotheses, generated models, and wrote the manuscript.

